# Sleep homeostasis in a naturalistic setting

**DOI:** 10.1101/2024.07.02.601682

**Authors:** Péter P. Ujma, Róbert Bódizs

**Affiliations:** Semmelweis University, Institute of Behavioural Sciences, Budapest, Hungary

**Keywords:** sleep homeostasis, synaptic homeostasis, delta wave, slow wave, NREM, sleep pressure, EEG, spectral exponents

## Abstract

Sleep, especially NREM sleep depth is homeostatically regulated, as sleep pressure builds up during wakefulness and diminishes during deep sleep. Previous evidence from this phenomenon, however, mainly stems from experimental studies which may not generalize to an ecologically valid setting. In the current study, we used a dataset of 246 individuals sleeping for at least seven nights each with a mobile EEG headband according to their ordinary daily schedule to investigate the effect of time spent in wakefulness on sleep characteristics. Increased time in wakefulness prior to sleep was associated with decreased sleep onset latency, increased sleep efficiency, a larger percentage of N3 sleep, and higher delta activity. Moreover, increased sleep pressure resulted in an increase in both the slope and the intercept of the sleep EEG spectrum. As predicted, PSD effects were most prominent in the earliest hours of sleep. Our results demonstrate that experimental findings showing increased sleep depth after extended wakefulness generalize to ecologically valid settings, and that time spent awake is an important determinant of sleep characteristics on the subsequent night. Our findings are evidence for the efficacy of sleep restriction, a behavioral technique already widely used in clinical settings, as a simple but powerful method to improve the objective quality of sleep in those with sleep problems.

## Introduction

Homeostasis is a key concept in understanding living systems. It covers the innate mechanisms of organisms that enable them to maintain steady internal states in a changing external environment (Modell et al., 2015). The concept of sleep homeostasis was coined to explain empirical evidence suggesting that several aspects of sleep depend on the recent sleep-wake history of the organism, thus sleep characteristics could be involved in maintaining the regulated balance between sleep and waking (Borbély and Tobler, 2023). A now widely received complete model of sleep characteristics is the two-process model (Borbély, 1982; Daan et al., 1984). In the two-process model, both homeostatic and circadian processes operate to regulate sleep and wakefulness. Homeostatic pressure for sleep builds up during wakefulness, while the phase of the circadian process determines the sleep threshold at which sleep is initiated or terminated. A prominent recent explanatory model, known as the synaptic homeostasis hypothesis, highlights the overall level of synaptic strengths within the central nervous system as the neurophysiological mechanism behind the two-process model (Tononi and Cirelli, 2014). In this model, sleep pressure reflects on the overall level of synaptic strengths within the central nervous system, whereas the function of sleep is to downscale these strengths. Based on this widely supported model, synaptic potentiation is an inevitable consequence of wakefulness, whereas deep sleep (and more specifically, the slow waves that occur in deep sleep) serves synaptic downscaling, enabled by low levels of noradrenergic activity.

Typical studies investigating the homeostatic regulation of sleep focus on recovery sleep after an experimentally scheduled extension of prior wakefulness (Borbély et al., 1981; Marzano et al., 2010) or the overnight dynamics of sleep as reflecting dissipating sleep pressure (Aeschbach and Borbély, 1993; G Horváth et al., 2022). Other commonly used research designs to unravel sleep homeostasis are the experimental extension of sleep opportunity by scheduled daytime naps (Campbell and Feinberg, 2005; Werth et al., 1996) or enforced bedrest (Klerman et al., 2021) and the non-24-hour day protocols known as forced desynchrony settings (Dijk and Czeisler, 1995). In each of these designs, changes in sleep metrics after the experimental manipulation of the duration of wakefulness are interpreted as evidence for sleep homeostasis.

Based on experimental research, several features of sleep macrostructure and electroencephalography (EEG) indeed exhibit finely graded homeostatic regulation. The propensity to initiate sleep (sleep latency) (De Gennaro et al., 2010; Rosenthal et al., 1993) total sleep time (TST) (Rosenthal et al., 1991), NREM sleep duration (Rosenthal et al., 1991), sleep efficiency (De Gennaro et al., 2010; Levine et al., 1988), slow wave sleep time and percentage (De Gennaro et al., 2010; Knowles et al., 1986; Webb and Agnew, 1971), and particularly slow wave activity (SWA, spectral power in the 0.75-4.5 Hz range) (Achermann et al., 1993) were shown to reflect at least in part the sleep-wake history of the organism. SWA is an indicator of sleep intensity that reflecting a restorative function of sleep (Borbély and Tobler, 2023). Recent evidence indicates a gradual levelling off of excess EEG power along the frequency scale, with lower frequencies showing stronger wake time-dependent upregulation. Consequently, sleep-wake history was best reflected by the steepness of the spectral slope of aperiodic activity in the sleep EEG (Bódizs et al., 2024).

Despite the strong focus on sleep homeostasis in academic research, facilitated by the decisive effect of the elegantly formulated two-process model of sleep regulation (Borbély, 1982; Daan et al., 1984), virtually all evidence for it stems from experimental studies. While experimental studies are powerful tools to demonstrate causality, they may suffer from deficiencies in ecological validity and their findings may not generalize to real-world settings (Holleman et al., 2020). In the specific case of sleep homeostasis, as recordings of undisturbed, natural sleep were virtually never analyzed to demonstrate this phenomenon, it is possible that the relatively drastic manipulations of wake time introduced in experimental studies are poor models of the much more modest day-to-day variability of sleep pressure, and in real-world settings sleep parameters are more robust to slight changes in the characteristics of the preceding wakefulness.

Here we aimed to fill this gap by analyzing the within-person covariation of wakefulness duration and subsequent sleep in a sample of healthy volunteers using wearable EEG-headbands on consecutive nights in an ecologically valid naturalistic setting. We focused on both macrostructural and spectral EEG indicators of sleep homeostasis to investigate if they reflect the characteristics of previous wakefulness. Our goal was to translate the concept of sleep homeostasis, conceived in laboratory-based intervention studies, to an ecologically valid setting and to demonstrate that small natural fluctuations in wake durations also influence sleep characteristics in line with the two-process model.

Our hypothesis was that sleep homeostasis is observable in the wild, that is, natural variations in the duration of wakefulness result in changes in subsequent sleep comparable to those observed in experimental studies. Specifically, we hypothesized that increased time in wakefulness results in 1) increased overall sleep propensity, shown by reduced sleep latency and sleep fragmentation as well as an increase in the relative proportion of deep sleep, 2) increased low-frequency activity in the sleep EEG as a marker of homeostatic restorative processes, especially in the delta frequency range and in the first hours of sleep, and 3) in line with the previous hypothesis, a steeper spectral slope.

## Methods

### Participants

We used data from the Budapest Sleep, Traits and Experiences Study (BSETS) a multiday observational study. The full protocol of BSETS has been published separately (Taji et al., 2023). In short, BSETS participants were health volunteers who tracked their life for at least seven consecutive days, including an array of demographic, psychological and psychiatric questionnaires, diaries about daily and nightly experiences, and an electroencephalographic (EEG) recording of each night sleep, in order to discover the relationship between sleep and daily experiences in a causally informative time-lagged within-participant design exploiting natural variation in daily experiences and sleep patterns. No experimental interventions or behavioral limitations were given, participants were free to schedule their daily activities and their sleep as they chose.

The current study relies on night EEG recordings. After discarding first-night recordings (see *Statistical analysis* ) 1365 nights with EEG recordings from 246 participants were available.

The Institutional Review Board (IRB) of Semmelweis University, as well as the Hungarian Medical Council (under 7040-7/2021/ EÜIG "Vonások és napi események hatása az alvási EEG-re" [“The effect of traits and daily activities and experiences on the sleep EEG”]), approved BSETS as compliant with the latest revision of the Declaration of Helsinki. All participants gave written informed consent on a form reviewed and approved by the IRB.

### Electroencephalography

BSETS participants were issued a Dreem2 mobile EEG headband (Arnal et al., 2020; Dreem Inc, 2017) and trained in its use to record their sleep at their homes for at least 7 consecutive nights. Dreem2 uses dry silicone electrodes to record brain activity with a 250 Hz sampling frequency, and a validated complimentary algorithm (Arnal et al., 2020; Dreem Inc, 2017) uses the resulting waveforms to score the vigilance state of participants. Based on these scorings, the following sleep macrostructure metrics were extracted: sleep efficiency (SE), total sleep time (TST), sleep onset latency (SOL), wake after sleep onset (WASO), the number of awakenings, and the latency, duration, and percentage of N1, N2, N3, and REM sleep. Sleep macrostructure metrics are often bounded at perfect values (e.g. SOL at 0 and SE at 100%), resulting in a skewed distribution. In order to make these variables more appropriate for linear analyses, we applied an outlier exclusion using the generalized form of Grubb’s test (implemented with the isoutlier() MATLAB function) (Pierson-Bartel and Ujma, 2024). We briefly report on findings without this data cleaning procedure (see Results).

For quantitative EEG (qEEG) analysis, we used the channel F7-O1 based on preliminary analyses (Taji et al., 2023) showing a good tradeoff between this channel’s data quality and its ability to record topographically widespread activity such as slow waves. For qEEG analyses, we used a complimentary algorithm to automatically detect artifactual EEG epochs on a 2-second basis. Based on preliminary analyses (Taji et al., 2023) and in line with published BSETS protocols (Pierson-Bartel and Ujma, 2024; Taji et al., 2023) epochs were discarded if this algorithm estimated artifact probability as greater than 25%. If the proportion of non-artifactual epochs for a night was below 20%, the night was discarded entirely. We used the periodogram() function in MATLAB EEGLab with 2-second nonoverlapping epochs and Hamming windows to perform spectral analysis. Power spectral density (PSD) estimates were averaged across epochs for each night and log10-transformed. Band-wise spectral values were calculated by averaging power values within the following frequency ranges: delta (0.5-4 Hz), theta (4-7 Hz), alpha (7-10 Hz), low sigma (10-13 Hz), high sigma (13-16 Hz), and beta (16-25 Hz) by averaging.

### Spectral parametrization

We decomposed absolute spectra into periodic and aperiodic components using FOOOF (“Fitting Oscillations & One Over f”, available at https://github.com/fooof-tools/fooof) (Donoghue et al., 2020). The power law function was estimated in the 3-20 Hz range. We allowed periodic components (spectral peaks) with a width of 1–4 Hz, a minimum peak height of 1 SD. We discarded nights where periodic and aperiodic components accounted for less than 95% of the variance in the power spectrum (N = 6). We used the spectral intercept and spectral slope as variables of interest indicating increased sleep pressure.

### Subjective sleep quality and dreaming

Upon awakening, participants reported their subjectively rated sleep quality using the Hungarian version of the Groningen Sleep Quality Scale (GSQS). Higher scores on this instrument indicate worse sleep. GSQS total were used as the main estimate of subjective sleep quality. In line with previous analyses (Pierson-Bartel and Ujma, 2024), we also specify two alternative metrics, the first unscored question of the GSQS (“I had a deep sleep last night”), treated as a binary variable, and an additional question prompting participants to rate their subjective level of restedness on a Likert scale of 1 to 10, which was treated as a continuous variable.

Participants also reported in this diary whether they recall a dream from the night. We used this variable to attempt to replicate previous research (De Gennaro et al., 2010) which indicated that dream recall is less frequent under increased sleep pressure.

### Time spent awake as the indicator of sleep pressure

Sleep pressure builds up steadily during wakefulness (Borbély and Tobler, 2023). Thus, we estimated the amount of sleep pressure at the beginning of the night as the time since the previous night’s last sleep epoch and the current night’s first sleep epoch (henceforth referred to as “time spent awake” or “time awake”). The clock time of both was based on EEG measurement.

EEG recordings were only performed at night, so sleep pressure may be overestimated if daytime naps are not considered. Participants recorded in their evening diary if they napped during the day and self-reported the total duration of napping. 297 daytime naps with an average duration of 70.34 minutes were recorded.

Because nap duration may be misreported and naps may be shallower than nighttime sleep, simply excluding self-reported naps from time awake may introduce bias. Therefore, in two alternative specification of the analyses we calculated the effect of time spent awake on sleep variables both with and without excluding naps.

### Statistical analysis

We investigated the relationship between sleep pressure and objective and subjective sleep metrics using multilevel models (McCrae et al., 2008; Robson and Pevalin, 2015), implemented using the fitglme() MATLAB function. Multilevel models are statistical tools which can handle dependent observations (such as multiple nights recorded from the same participant) and can estimate if the effect of a predictor exists at the within-person level (Level 1) or the between-person level (Level 2), also allowing controlling for covariates at both levels. For example, a within-person effect exists if the same person has deeper sleep on nights with greater sleep pressure, while a between-person exists if participants who have higher average sleep pressure across nights also have deeper sleep on average. Within-person effects are more in line with a causal interpretation as they reflect that the same person’s sleep changes after increased sleep pressure, as opposed to between-person effects which merely reflect a correlation observed between participants (Pierson-Bartel and Ujma, 2024).

We modeled the effect of sleep pressure by first creating a between-person and a within-person specification of sleep pressure (McCrae et al., 2008; Robson and Pevalin, 2015). For the between-person specification, the mean amount of time spent awake (across all days) was entered for all observations from the same person. For the within-person specification, deviations from the individual mean were used, with each person’s data centered around 0. A separate model was fitted for each sleep macrostructure variable, EEG power band, and subjective sleep metric as dependent variables. For binwise qEEG analyses, a separate model fitted for each frequency bin. Each model used a random intercept per participant, both the between-person and the within-person specification of sleep pressure as predictors, together with age, sex, a dummy variable for weekend/weekday, and lagged outcomes (the previous day’s value of the dependent variable) to control models for these potential confounders.

From each model, the critical result was the within-person regression coefficient: the estimated causal effect of an additional hour of time spent awake on a given person’s sleep. The p-values of these regression coefficients were corrected for multiple testing using the Benjamini-Hochberg method of false discovery rate correction (Benjamini and Hochberg, 1995). All nominally significant (p<0.05) p-values reported in the manuscript also survive this correction unless indicated otherwise.

Because time spent awake (calculated since the end of the first night’s sleep) could only be estimated starting on the second night, first night recordings could not be directly used for our analyses. We did use first night recordings, however, to control for lagged outcomes to eliminate spillover effects from the previous nights. For example, a model estimating the effect of time spent awake on WASO is controlled for WASO on the previous night. Controls for lagged outcomes was another separate reason for not using first-night recordings.

### Data availability

Raw data is available at https://osf.io/2p8hj/.

## Results

### Descriptive statistics

The mean age of the sample was 29.15 years (SD=12.82), with 55% females and 45% males. The mean time spent awake was 17.05 hours (SD=2.03). Intraclass coefficients ranged from 0.076 (time spent awake) to 0.718 (alpha power). Detailed descriptive statistics (valid Ns, means, SDs, intraclass correlation coefficients and within- and between-participant correlations are available in the **Supplementary file 1** .

### Sleep macrostructure

Increased time spent awake during the previous day was associated with deeper and less disturbed sleep (**Figure 1** ). For each additional hour spent awake, sleep onset latency was reduced by 0.62 minutes and wake after sleep onset by 1.31 minutes, sleep efficiency was increased by 0.23 percentage points and 0.5 fewer awakenings were observed.

**Figure 1.**
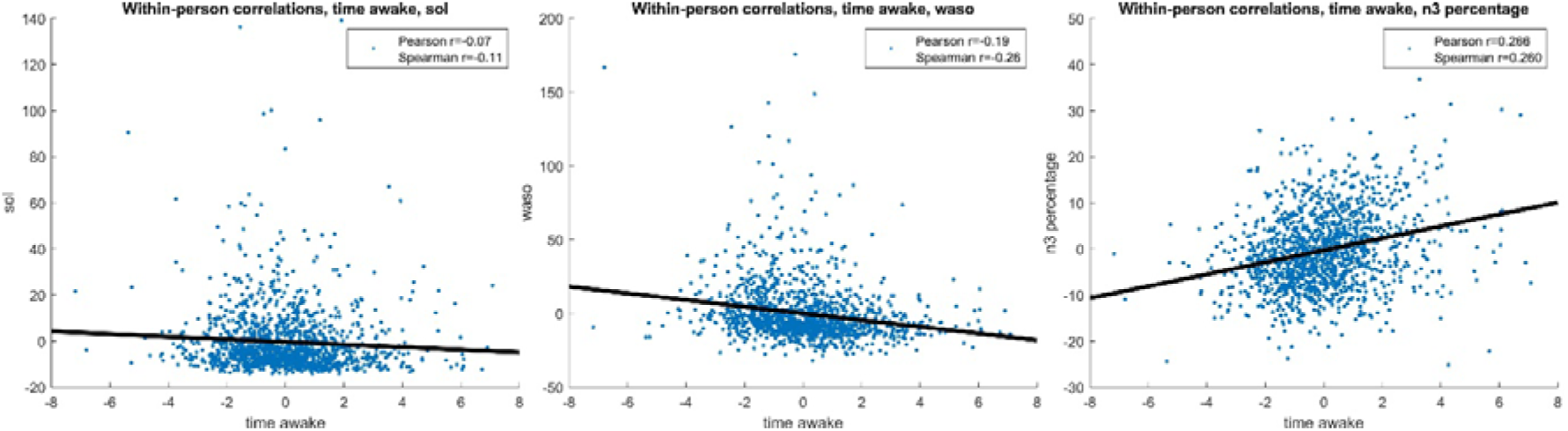
Selected effects of increased sleep pressure. The scatterplots show the within-participant relationship between time spent awake (horizontal axis) and sleep onset latency (left), wake after sleep onset (middle) and N3 percentage (right). All values are deviations from the individual mean, hence they are centered around zero. All variables are residualized to age, sex and day of the week. Pearson and Spearman correlations are shown.

Total sleep time was reduced after increased sleep pressure: however, this is expected as longer wakefulness results in less time to sleep due to the normal obligations of the following day. Consequently, increased sleep pressure was associated with later bedtimes (32 minutes for each additional hour spent awake). Controlling for bedtime reduced the association between sleep pressure and total sleep time by 70% to 4.13 minutes, but it remained statistically significant (p=0.002).

The analysis of sleep composition revealed that – mirroring reductions in total sleep time – time spent in N1, N2 and REM was significantly reduced after longer wakefulness, but N3 duration was not significantly changed. When we used percentages instead of total time, the percentage of N3 sleep was increased, N2 sleep was reduced, while N1 and REM were unchanged.

All effects replicated with only slight variation in the effect sizes after naps were excluded (**Table 1** ).

**Table 1.** Within-person effects of time spent awake on sleep macrostructure. In each line, we show the expected effect of one additional hour spent awake during the previous day on the sleep macrostructure indicators in the first column (B), the standard error and the p-value of this coefficient. Values are expressed in natural units (in parentheses). All models are controlled for age, sex and day of the week (weekday/weekend). The left half of the table shows effects without and the right half with the inclusion of self-reported naps from time spent awake.

|  | Naps included |  |  | Naps excluded |  |  |
| --- | --- | --- | --- | --- | --- | --- |
|  | B | SE | p | B | SE | p |
| Sleep onset latency (min) | -0.616 | 0.217 | 0.005 | -0.633 | 0.213 | 0.003 |
| Sleep efficiency (%) | 0.225 | 0.097 | 0.021 | 0.268 | 0.095 | 0.005 |
| Awakenings (count) | -0.488 | 0.124 | <0.001 | -0.535 | 0.119 | <0.001 |
| WASO (min) | -1.313 | 0.299 | <0.001 | -1.287 | 0.283 | <0.001 |
| Total sleep time (min) | -13.518 | 1.394 | <0.001 | -11.005 | 1.404 | <0.001 |
| **N1 duration (min)** | -0.619 | 0.141 | <0.001 | -0.637 | 0.135 | <0.001 |
| **N2 duration (min)** | -9.122 | 0.853 | <0.001 | -7.764 | 0.837 | <0.001 |
| **N3 duration (min)** | -0.325 | 0.380 | 0.392 | -0.008 | 0.376 | 0.983 |
| **REM duration (min)** | -1.823 | 0.625 | 0.004 | -1.818 | 0.614 | 0.003 |
| **N1 percentage (%)** | 0.010 | 0.031 | 0.755 | -0.018 | 0.030 | 0.541 |
| **N2 percentage (%)** | -0.785 | 0.120 | <0.001 | -0.686 | 0.117 | <0.001 |
| **N3 percentage (%)** | 0.630 | 0.122 | <0.001 | 0.647 | 0.119 | <0.001 |
| **REM percentage (%)** | 0.181 | 0.115 | 0.117 | 0.088 | 0.113 | 0.438 |

### Quantitative EEG: Power spectral density

Increased time spent awake led to NREM sleep EEG PSD increases in the delta through the low sigma frequency range (**Table 2** ).

**Table 2.** Within-person effects of time spent awake on absolute spectral power of the NREM sleep EEG. In each line, we show the expected effect of one additional hour spent awake during the previous day on the power spectral density (PSD) of the EEG band indicated in the first column (B), the standard error (SE) and the p-value of this coefficient. PSD values were log10-transformed before analyses. All models are controlled for age, sex, and day of the week (weekday/weekend). The left half of the table shows effects without and the right half with the inclusion of self-reported naps from time spent awake. ± indicates a nominally significant (p<0.05) effect which does not survive correction for multiple testing.

|  | Naps included |  |  | Naps excluded |  |  |
| --- | --- | --- | --- | --- | --- | --- |
|  | B | SE | p | B | SE | p |
| **Delta** | 0.011 | 0.003 | <0.001 | 0.010 | 0.003 | <0.001 |
| **Theta** | 0.008 | 0.002 | <0.001 | 0.008 | 0.002 | <0.001 |
| **Alpha** | 0.008 | 0.002 | <0.001 | 0.008 | 0.002 | <0.001 |
| **Low sigma** | 0.005 | 0.002 | 0.002 | 0.004 | 0.002 | 0.016 |
| **High sigma** | 0.003 | 0.002 | 0.047<sup>±</sup> | 0.002 | 0.002 | 0.208 |
| **Beta** | 0.003 | 0.002 | 0.199 | 0.002 | 0.002 | 0.262 |

In more detailed analyses, we calculated a multilevel model for each 0.5 Hz frequency bin. **Figure 2** illustrates the expected change in the PSD of each frequency bin per one of our additional wakefulness. Increased PSD was most prominent in the ∼1-2 Hz frequency range corresponding to slow waves, but the PSD increase remained at least nominally significant into the high alpha-low sigma frequency ranges.

**Figure 2.**
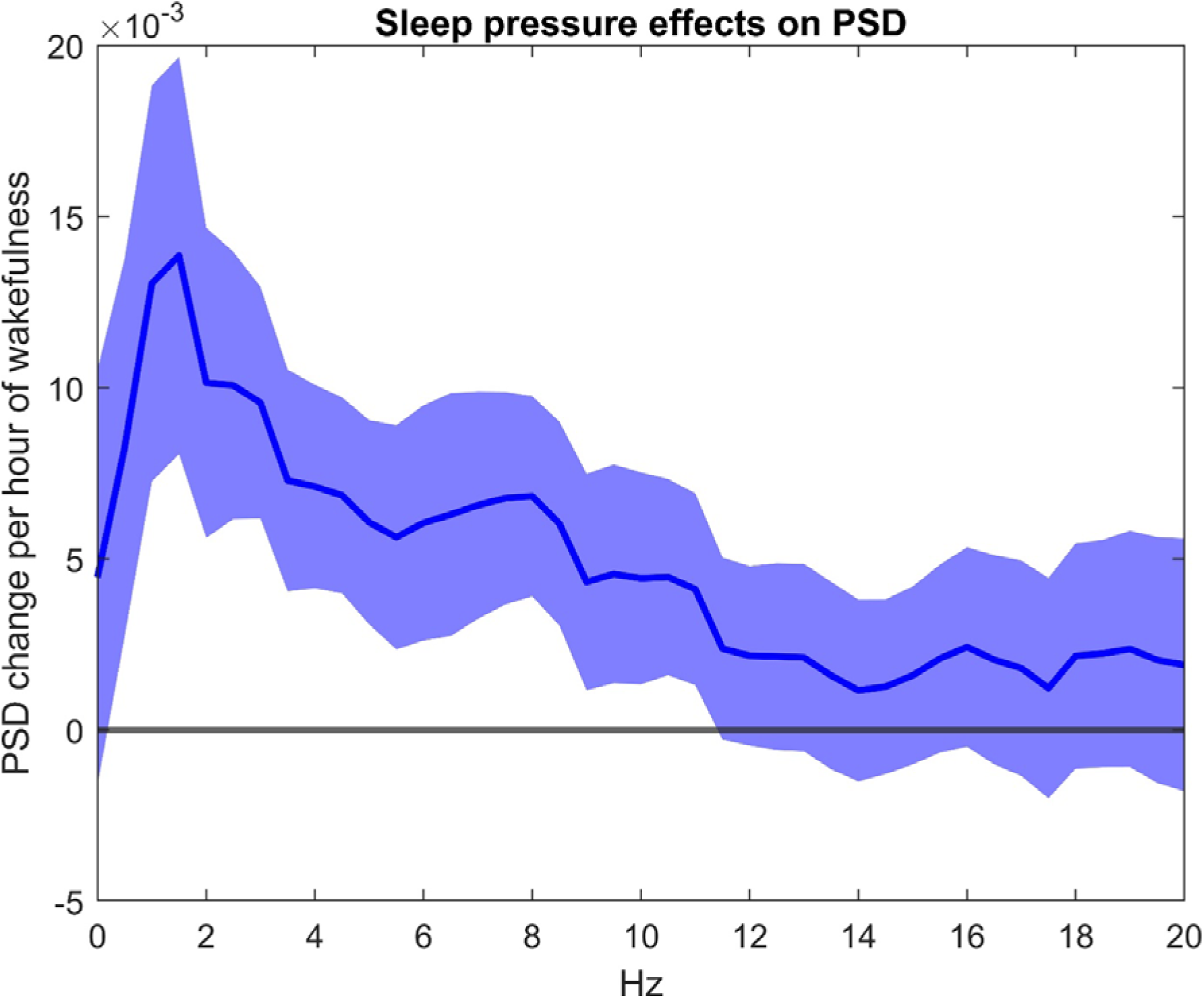
Within-person effects of time spent awake on binwise NREM sleep EEG PSD. The chart shows the within-person estimates of the effect of one additional hour spent awake during the previous way on the NREM sleep EEG PSD of each frequency. PSD values were log10-transformed before analyses. All models are controlled for age, sex, and day of the week (weekday/weekend). The shaded area shows 95% confidence intervals.

### Quantitative EEG: bi-hourly PSD

In the next step, we calculated NREM PSD separately from bi-hourly periods (with one hour of overlap up to the fifth hour) of recordings to test how homeostatic effects are distributed across the night.

As the duration of recordings and the distribution of artifact-free NREM sleep was non-uniform across the nights, the total number of available EEG nights varied by hour. We limited analyses for the first five hours of recordings in two-hour overlapping batches. For these batches, a total of 1632, 1606, and 1508 nights (including first nights) were available. Statistical modelling was performed in a manner identical to full-night recordings.

For all periods, a trend for increased low-frequency activity as a function of increased time spent awake was observed. This trend, however, only reached significance across a broad frequency range in the first two hours of sleep. After the first two hours of sleep, significance was limited to a single isolated bin in the theta ranges.

The results are summarized in **Figure 3** .

**Figure 3.**
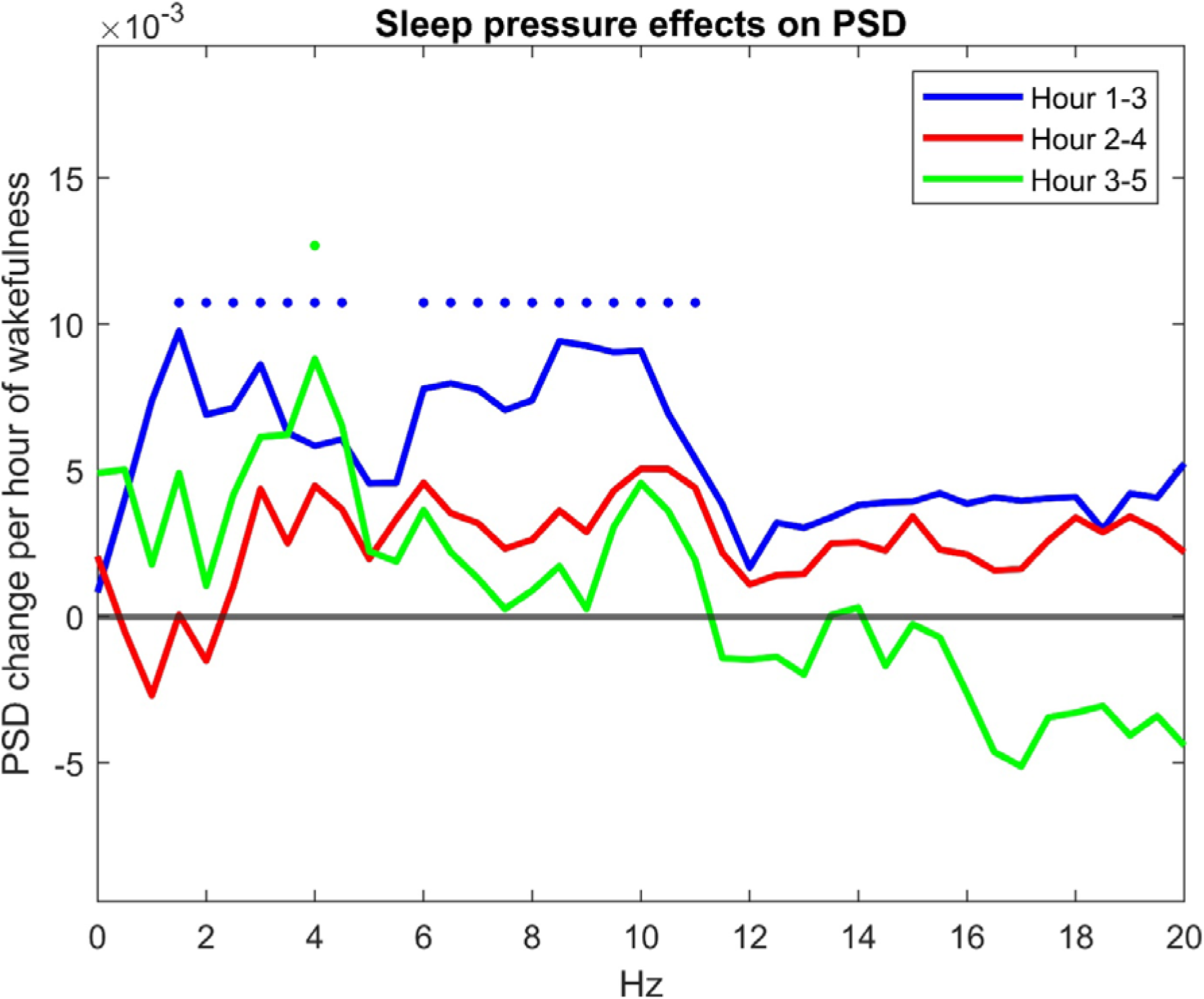
Within-person effects of time spent awake on binwise NREM sleep EEG PSD. The chart shows the within-person estimates of the effect of one additional hour spent awake during the previous way on the NREM sleep EEG PSD of each frequency, calculated separately for three bi-hourly, overlapping period of recordings (separate lines). PSD values were log10-transformed before analyses. All models are controlled for age, sex, and day of the week (weekday/weekend). Dots above the lines indicate that the effect is significant after correcting for multiple comparisons in the hour indicated by the line with the matching color.

### Quantitative EEG: spectral slopes and intercepts

We found that both spectral slopes and intercepts were significantly increased after increased time spent awake, mirroring results about PSD (**Table 3** ).

**Table 3.** Within-person effects of time spent awake on aperiodic spectral parameters. In each line, we show the expected effect of one additional hour spent awake during the previous day on the spectral intercepts and slopes, as indicated in the first column (B), the standard error and the p-value of this coefficient. All models are controlled for age, sex, and day of the week (weekday/weekend). The left half of the table shows effects without and the right half with the inclusion of self-reported naps from time spent awake.

|  | Naps included |  |  | Naps excluded |  |  |
| --- | --- | --- | --- | --- | --- | --- |
|  | B | SE | p | B | SE | p |
| **Intercept** | 0.014 | 0.004 | <0.001 | 0.013 | 0.004 | <0.001 |
| **Slope** | 0.009 | 0.004 | 0.018 | 0.009 | 0.004 | 0.011 |

Figure 4 shows the predicted spectral intercept and slope across a range of plausible wakefulness durations.

**Figure 4.**
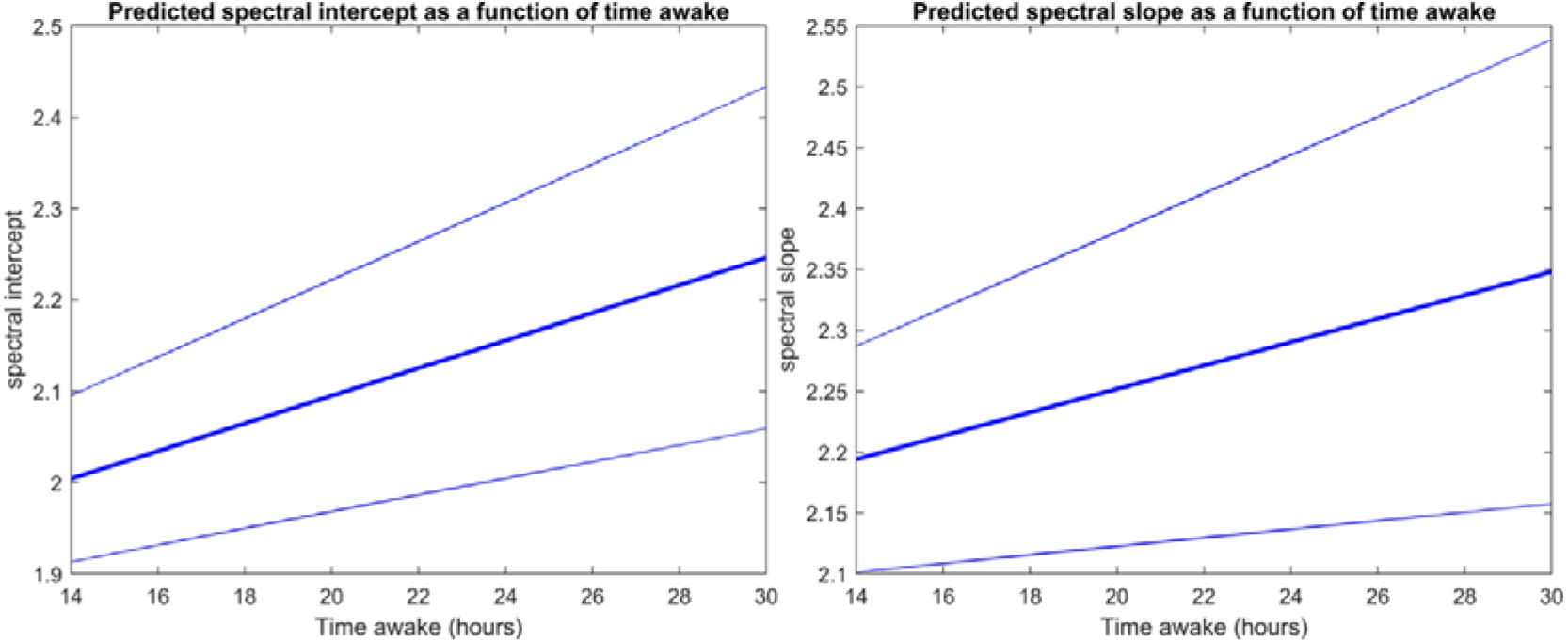
Predicted values of the spectral intercept (left) and spectral slope (right) as a function of the duration of previous wakefulness.

### Subjective sleep quality

We found no significant effect of time spent awake on self-reported subjective sleep quality, either considering GSQS total scores, the first item of the GSQS not counted towards the total score, or self-reported restedness on a Likert scale (**Table 4** ). Thus, although increased sleep pressure resulted in objectively deeper and more efficient sleep, this was not translated into a perception of higher sleep quality the following morning.

**Table 4.** Within-person effects of time spent awake on three measures of self-reported sleep quality: GSQS total score, the first unscored GSQS item, and self-reported restedness. (For the first GSQS item, the linear effect is shown for simplicity, although this estimate is derived from logistic regression as this is a binary variable). In each line, we show the expected effect of one additional hour spent awake during the previous day on the sleep quality indicators in the first column (B), the standard error and the p-value of this coefficient. Values are expressed in raw scores. All models are controlled for age, sex, and day of the week (weekday/weekend). The left half of the table shows effects without and the right half with the inclusion of self-reported naps from time spent awake.

|  | Naps included |  |  | Naps excluded |  |  |
| --- | --- | --- | --- | --- | --- | --- |
|  | B | SE | p | B | SE | p |
| **GSQS** | 0.020 | 0.057 | 0.722 | 0.011 | 0.055 | 0.847 |
| **GSQS1** | -0.009 | 0.048 | 0.854 | 0.003 | 0.046 | 0.941 |
| **Restedness** | 0.010 | 0.038 | 0.788 | -0.019 | 0.037 | 0.611 |

### Dream recall

We found no compelling evidence that increased sleep pressure reduces the chance of reporting a dream in the morning diary. While the effect sizes were in this direction both with (within-participant OR=0.94 per hour of wakefulness, p=0.12) and without (OR=0.95, p=0.11) the exclusion of naps, they were not statistically significant.

### Between-participant effects

Our research focused on within-participant effects, that it, how a given participant’s sleep changed after varying degrees of wakefulness. However, between-participant effects (associations between average wake duration and average sleep characteristics across participants) are also of interest.

Between-participant effects of wakefulness were very similar to within-participant effects, although less precisely estimated due to fewer observations (Figure 5 ).

**Figure 5.**
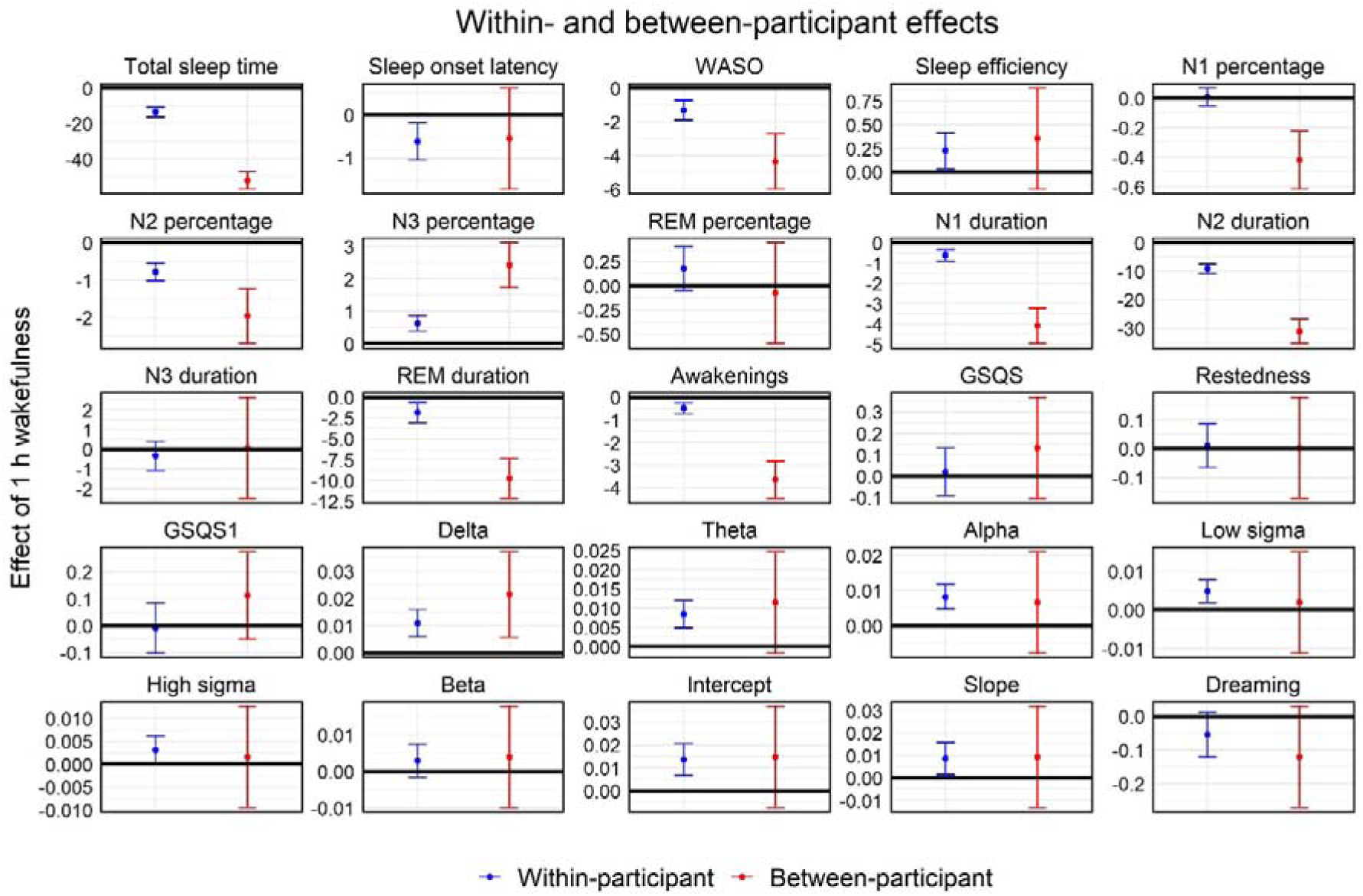
Comparison of within-and between participants effects. Each tile shows the estimated within- and between-participant effects of one hour of wakefulness (without taking into account naps) on sleep characteristics. Estimates originate from multilevel models adjusted for age, sex and day of the week. Error bars show 95% confidence intervals.

The duration of sleep and specific sleep stages was much more strongly associated with wakefulness, which is unsurprising as these are zero-sum variables at the between-participant level (on average, a given participant spends a certain amount of time sleeping and the rest awake). However, longer typical wakefulness was also associated with significantly higher delta power, N3 percentage, significantly reduced WASO and awakenings, and a within-participant-like trend for lower sleep onset latency, higher sleep efficiency, higher spectral slopes and intercepts, and higher EEG power even at higher frequency ranges. Notably, despite the zero-sum nature of sleep and wakefulness, typical N3 duration was not affected by typical wake duration, suggesting a compensatory mechanism where the lower absolute amount of sleep is composed of deeper stages. These findings show that homeostatic processes operate on the between-participant level as well, and – even after controlling for age – participants with longer typical wakefulness tend to have increased sleep propensity.

### Outlier effects

We elected to exclude observations with extreme values to make analyses more amenable for linear analyses (see Methods). In **Supplementary file 2** , we report regression results without this correction. Outlier exclusion made minimal effect on the results, and except for affecting marginal significance in the case of two between-participant effects it was limited to changes in the signs of otherwise non-significant findings.

## Discussion

The two-process model represents a powerful explanatory framework for homeostatic sleep regulation. Evidence for this model, however, has previously only been sourced from experimental studies. This is potentially problematic as experiments in the biomedical sciences often fail to translate into ecologically valid settings (Drude et al., 2021). In the current case, it is possible that while drastic experimental manipulations in wake duration can affect sleep characteristics, the effect of much smaller variations in wake duration experienced by a typical person are overpowered by other determinants of sleep, such as stress (Åkerstedt et al., 2012; Yap et al., 2022), learning or cognitive load (Cerasuolo et al., 2019), auditory stimulation (Cantero et al., 2002), or meals (Duan et al., 2021; St-Onge et al., 2016). In the current study, we show that experimental studies of sleep homeostasis indeed replicate in an ecologically valid, naturalistic study, and sleep characteristics change as a function of sleep-wake history within the normal range.

Longer wakefulness resulted in an increased propensity for sleep, especially of deep sleep and slow waves. Specifically, longer pre-sleep wakefulness was followed by reduced sleep onset latency, increased sleep efficiency and higher N3 percentage at the expense of other sleep stages, as well as decreased WASO and fewer awakenings. Band- and binwise spectral analysis replicated experimental laboratory studies (Aeschbach and Borbély, 1993; Borbély et al., 1981; Marzano et al., 2010) and reveal that increased wakefulness results in increased NREM sleep PSD, most prominently at the lowest frequencies but extending up to the sigma range. Binwise statistics clearly revealed higher frequencies were progressively less involved in sleep homeostasis (Bódizs et al., 2024): effects transcend the canonical SWA range, extend over higher frequencies, but increasing EEG frequencies are characterized by decreasing sensitivity to extensions in pre-sleep wakefulness. Moreover, both the NREM sleep EEG spectral slope and exponents values increased as a function of time awake, supporting the relevance of these indices in depicting sleep homeostasis.

Importantly, we found no evidence that increased wakefulness duration also resulted in improvements in subjectively rated sleep quality the next morning. We also could not replicate previous finding (De Gennaro et al., 2010) that increased sleep pressure results in a reduced likelihood of dreaming.

Our findings provide strong empirical evidence that homeostatic mechanisms regulate sleep even under the naturalistic conditions of daily life. In our ecologically valid observational study, as in previous laboratory studies (Aeschbach and Borbély, 1993; Borbély et al., 1981; Marzano et al., 2010), sleep pressure increases with time spent awake and translated into an increased propensity for sleep initiation.

Sleep characteristics do not vary randomly from night to night, but to some extent they are habitual or trait-like (Tucker et al., 2007). This was also observed in our dataset (**Supplementary file 1** ): on average, 41% of the variance of macrostructure variables, 66% of PSD values, 57% of spectral parameters, and 21% of subjective sleep quality reports was at the within-participant level, showing that the same participant experiences similar sleep across nights. Between-participant analyses using habitual sleep metrics generally replicated the within-participant effects of time awake on sleep propensity and structure. Participants with habitually longer periods of wakefulness were characterized by higher N3 percentage, delta power, as well as by significantly reduced WASO and number of awakenings, with trends in line with within-participant effects for most other sleep metrics as well. These findings suggest that while fluctuations in a person’s sleep propensity can reflect fluctuations sleep pressure, habitual sleep patterns may also be partially the result of the habitual duration and intensity of wakefulness.

Our results have important implications for both sleep research and somnology. The replication of laboratory studies of sleep homeostasis provides not only evidence for the two-process model, but also for the feasibility of conducting sleep research outside of the laboratory. Using wearable EEG devices at home is substantially less cost and labor intensive than standard laboratory PSG studies and this approach may result in larger, better-powered studies to test hypotheses about sleep regulation and function with relatively little compromise towards fidelity. Studying impairments of sleep regulation in a naturalistic setting might especially be useful to understand the mechanisms beyond insomnia, depression, and circadian rhythm disorders.

A special case where sleep homeostasis under natural conditions is important is sleep restriction therapy (SRT), commonly used to alleviate symptoms of insomnia. In SRT, patients are instructed to reduce time spent asleep (or attempting to sleep) to increase sleep pressure and the depth and efficiency of subsequent sleep. A recent meta-analysis of SRT RCTs (Maurer et al., 2021) supported the efficacy of SRT to reduce the insomnia severity index, sleep onset latency and WASO, but this evidence was rated as low quality due to the low number of studies, between-study heterogeneity, and lack of long-term follow-up. Our study using a causally informative non-RCT design confirms that SRT might be a promising therapy for sleep problems, and that increasing time spent awake may be a highly efficacious way to increase sleep quality. Caution must be warranted, however, as SRT was shown to negatively affect vigilance and sleepiness (Kyle et al., 2014) suggesting that while sleep following SRT is biologically more efficient, its restorative properties might be missing. In line with these observations, we also found that next-morning subjective sleep quality was not significantly improved after longer time spent awake (and the trend was for a negative change). Wearable instruments, as presented in the current study, could be used in clinical settings to fine-tune SRT in way that balances the efficiency and restorative property of sleep in individual patients.

Our work has limitations. First, ours was a relatively young volunteer sample without clinically significant sleep complaints or other medical issues. The findings may not generalize to older populations or those with significant sleep problems, requiring replication. Second, our study relied on a mobile EEG headband instead of gold-standard PSG to measure sleep. While the general reliability of the method has been demonstrated in previous studies, sleep structure may be better determined using PSG. This limitation especially affects qEEG findings, as the Dreem2 headband has strong hardware filtering above 18 Hz and the accurate mapping of higher frequencies is not possible with this method. Third, we could imperfectly model events in wakefulness which may affect the amount of sleep pressure that was generated. While we had self-reported data on naps which did not substantially affect findings, an ideal study would include actigraphic measures of the day which could precisely model sleep pressure by taking into account the exact duration of napping and physical activity.

In sum, our findings demonstrate that validity of the two-process model in the description of day-to-day natural variations in sleep. Similarly to experimental manipulations, a naturally longer wakefulness period also results in increased sleep propensity and depth but no change in subjective sleep quality, informing clinical interventions aimed to improve sleep.

## Acknowledgements

This was supported by the National Research, Development and Innovation Office – NKFIH (grant number: 138935). This research has been implemented with the support provided by the Ministry of Innovation and Technology of Hungary from the National Research, Development and Innovation Fund, financed under the TKP2021-EGA-25 funding scheme. Research support was also provided by the Culture and Innovation Ministry of Hungary from the National Research, Development and Innovation Fund, financed under the TKP2021-NKTA funding scheme (project nr.: TKP2021-NKTA-47).

